# Determination of essential phenotypic elements of clusters in high-dimensional entities - DEPECHE

**DOI:** 10.1101/396135

**Authors:** Axel Theorell, Yenan Troi Bryceson, Jakob Theorell

## Abstract

Technological advances have facilitated an exponential increase in the amount of information that can be derived from single cells, necessitating new computational tools that can make this highly complex data interpretable. Here, we introduce DEPECHE, a rapid, parameter free, sparse k-means-based algorithm for clustering of multi-and megavariate single-cell data. In a number of computational benchmarks aimed at evaluating the capacity to form biologically relevant clusters, including flow/mass-cytometry and single cell RNA sequencing data sets with manually curated gold standard solutions, DEPECHE clusters as well or better as the best performing state-of-the-art clustering algorithms. However, the main advantage of DEPECHE, compared to the state-of-the-art, is its unique ability to enhance interpretability of the formed clusters, in that it only retains variables relevant for cluster separation, thereby facilitating computational efficient analyses as well as understanding of complex datasets. An open source R implementation of DEPECHE is available at https://github.com/theorell/DepecheR.

**Author summary:** DEPECHE-a data-mining algorithm for mega-variate data

Modern experimental technologies facilitate an array of single cells measurements, *e.g.* at the RNA-level, generating enormous datasets with thousands of annotated biological markers for each of thousands of cells. To analyze such datasets, researchers routinely apply automated or semi-automated techniques to order the cells into biologically relevant groups. However, even after such groups have been generated, it is often difficult to interpret the biological meaning of these groups since the definition of each group often dependends on thousands of biological markers. Therefore, in this article, we introduce DEPECHE, an algorithm designed to simultaneously group cells and enhance interpretability of the formed groups. DEPECHE defines groups only with respect to biological markers that contribute significantly to differentiate the cells in the group from the rest of the cells, yielding more succinct group definitions. Using the open source R software DepecheR on RNA sequencing data and mass cytometry data, the number of defining markers were reduced up to 1000-fold, thereby increasing interpretability vastly, while maintaining or improving the biological relevance of the groups formed compared to state-of-the-art algorithms.

## Introduction

Since the introduction of the first single colour flow cytometers in the 1960s, there has been a remarkable increase in the complexity of data that can be generated with single-cell resolution. Currently, flow and mass cytometers able to simultaneously assess up to 40 cellular traits are becoming widely available [1]. In parallel, the development of high-throughput sequencing technology has facilitated deep single-cell transcriptomic analyses [2]. Furthermore, development of high-resolution single-cell proteomic analyses are underway [3].

These technological advances necessitate new computational approaches to analyses of multi-and megavariate single cell data [4–7]. Previous algorithms have contributed to automating analyses, thereby enhancing reproducibility and avoiding a need for *a priori* biological knowledge for design of manual gating analysis strategies. Automated analysis algorithms, not restricted to uni-or bivariate displays of the data, have also made it possible to display much more of the information embedded in multivariate data. To date, however, manual gating strategies are still dominantly used, which in part is likely due to that it is easy to interpret what population a certain gate refers to, as it is defined by few markers. In an attempt to combine the objectiveness and reproducibility of automated analysis pipelines with the high interpretability of manual gating strategies, we have developed an algorithm termed Determination of Essential Phenotypic Elements of Clusters in High-dimensional Entities (DEPECHE). DEPECHE simultaneously clusters and simplifies the data by identifying the variables that contribute to separate individual clusters from the rest of the data. We have implemented DEPECHE in R (in the open source package DepecheR), providing a complete software suite for statistical analysis and visualization of single cell omics data.

## Results and Discussion

DEPECHE uses a penalized k-means clustering algorithm, related to the standard k-means algorithm [8]. In penalized k-means, a penalty term is introduced to the clustering algorithm. The value of the penalty, *λ*, determines the clustering resolution. Low clustering resolution implies that few clusters defined by few variables (high sparsity) are produced, and vice versa [9] (see online methods). Note that if *k* is high enough not to be limiting, the resolution of the emerging clusters depends entirely on the magnitude of *λ* and not on k, since DEPECHE annihilates all clusters that are pulled to the origin by the penalty. In DEPECHE, the penalty *λ* is tuned to identify the most *reproducible* clustering resolution, here termed the “optimal resolution”[10]. To illustrate what we mean by reproducible, we constructed a show case, featuring a bi-variate dataset *D* (Fig 1a). Visually, the dataset *D* contains three clusters, where the centers of the two larger clusters are located close to either axis. For these two clusters, one variable is sufficient to define their position. Now, if multiple datasets were generated from the same data source as *D*, for example by repeated experiments, we assume that they would contain the same clusters. Hence, imposing the optimal penalty *λ*_*i*_ (that corresponds to the optimal resolution of *D*) on all these datasets should ideally always result in the same clusters (high reproducibility). Contrarily, when clustering the same datasets with a penalty *λ* that differ significantly from the optimal penalty, the stochastic differences between the datasets are likely to induce solutions that deviate in cluster number, number of defining variables, and cluster center positions. In practice, DEPECHE tests a range of penalty values (*λ*_1_ < ⋯ < *λ*_*n*_), each on a collection of dataset pairs which are generated by sampling *N*_*R*_ data points from *D* (Fig 1b) with resampling. The optimal resolution is defined as the penalty *λ*_*i*_, which yields the lowest average variability within each dataset pair, as measured by the Adjusted Rand Index (ARI) [11]. In our example, this corresponds to the penalty *λ*_*i*_ that yields 3 clusters, since 3 similar clusters are identified in each resampled dataset of *D* (Fig 1c). The penalties *λ*_1_ and *λ*_*n*_ are considered suboptimal, since with these penalties, the stochastic differences in the resampled datasets lead to less coherent clustering results compared to results obtained with the optimal penalty *λ*_*i*_. From here (Fig 1c) DEPECHE uses two alternative routes. If the number of data points in the dataset *D* is high (as default > 10^4^), the most generalizable cluster centers that were produced using the optimal penalty are chosen (see online methods) and the data points of *D* are allocated directly to their closest cluster center (Fig 1d). If the dataset *D* has few data points, the full dataset *D* is clustered using the optimal penalty *λ*_*i*_ (Fig 1e).

**Fig 1:**
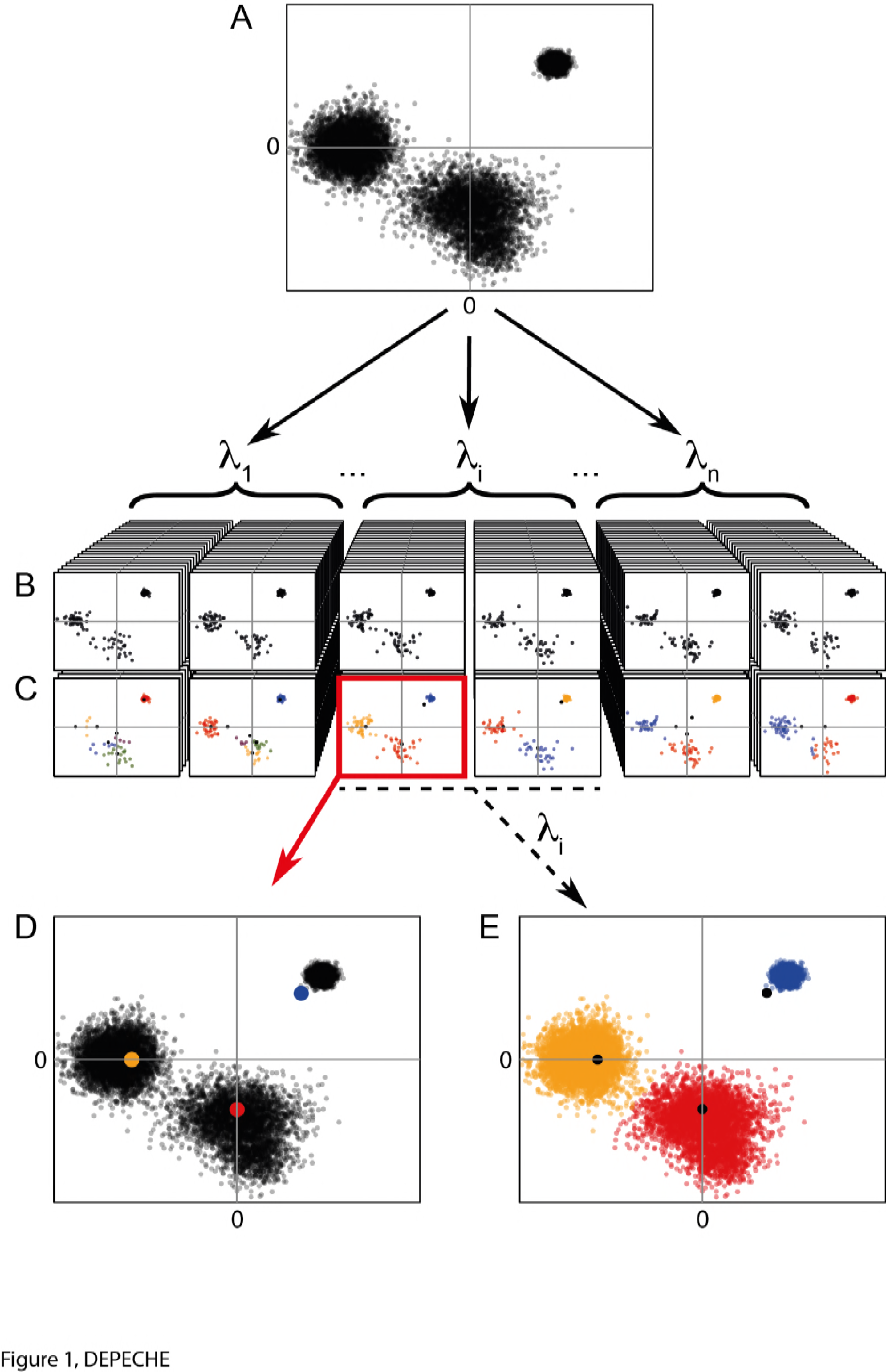
Illustration of the DEPECHE workflow: a) The original dataset *D*. b) *n* resampled datasets with *N*_*R*_ data points per dataset, are generated by sampling data points from *D* with resampling. Each resampled dataset has a corresponding penalty *λ*_*i*_ (*i =* 1,,, *n*). c) Each dataset in b is clustered with sparse k-means, using its corresponding penalty *λ*_*i*_. The red frame highlights the clustering with the strongest attractors, *i.e.* the most generalizable solution (see online methods). d, e) Finally, the full dataset is clustered by allocating each data point to its closest cluster center, using the most generalizable cluster center solution produced in b.

To evaluate how biologically accurate DEPECHE clustering is on mass cytometry data, a 32-variate mass cytometry bone marrow dataset [12] was clustered, and the overlap to 14 manually pre-defined cell populations was quantified using the ARI. With this dataset, DEPECHE identified 7 clusters at the optimal resolution, corresponding to all large pre-defined cell populations and to agglomerates of smaller cell populations, rendering an average ARI of 0.96, where an ARI of 1 corresponds to exact reproduction and an ARI of 0 means that the produced clusters are no more accurate than random allocation. (Fig 2a-b, S Fig 1a). Furthermore, using DEPECHE, the number of variables defining each cluster was reduced from 32 to a range from 8 to 28, thereby enhancing interpretability (Fig 2d). When comparing to other state-of-the-art clustering algorithms [12–17], DEPECHE obtained similar ARI as the best algorithms for both the 32-variate dataset and another 14-variate, 24 population, mass cytometry dataset [18] (S Fig 2a, Table 1).

**Table 1:** background information on all datasets

| Dataset | Data origin | n cells | n variables in analysis | n clusters in original |
| --- | --- | --- | --- | --- |
| Levine | Mass cytometry | 104184 | 32 | 14 |
| Bendall | Mass cytometry | 81747 | 14 | 24 |
| Björklund | scRNAseq | 648 | 35177 | 4 |
| Biase | scRNAseq | 56 | 19571 | 3 |
| Deng | scRNAseq | 268 | 13867 | 10 |
| Goolam | scRNAseq | 124 | 15487 | 5 |
| Kolodziejczyk | scRNAseq | 704 | 15117 | 3 |
| Pollen | scRNAseq | 301 | 13860 | 11 |
| Yan | scRNAseq | 90 | 13608 | 7 |

**Fig 2:**
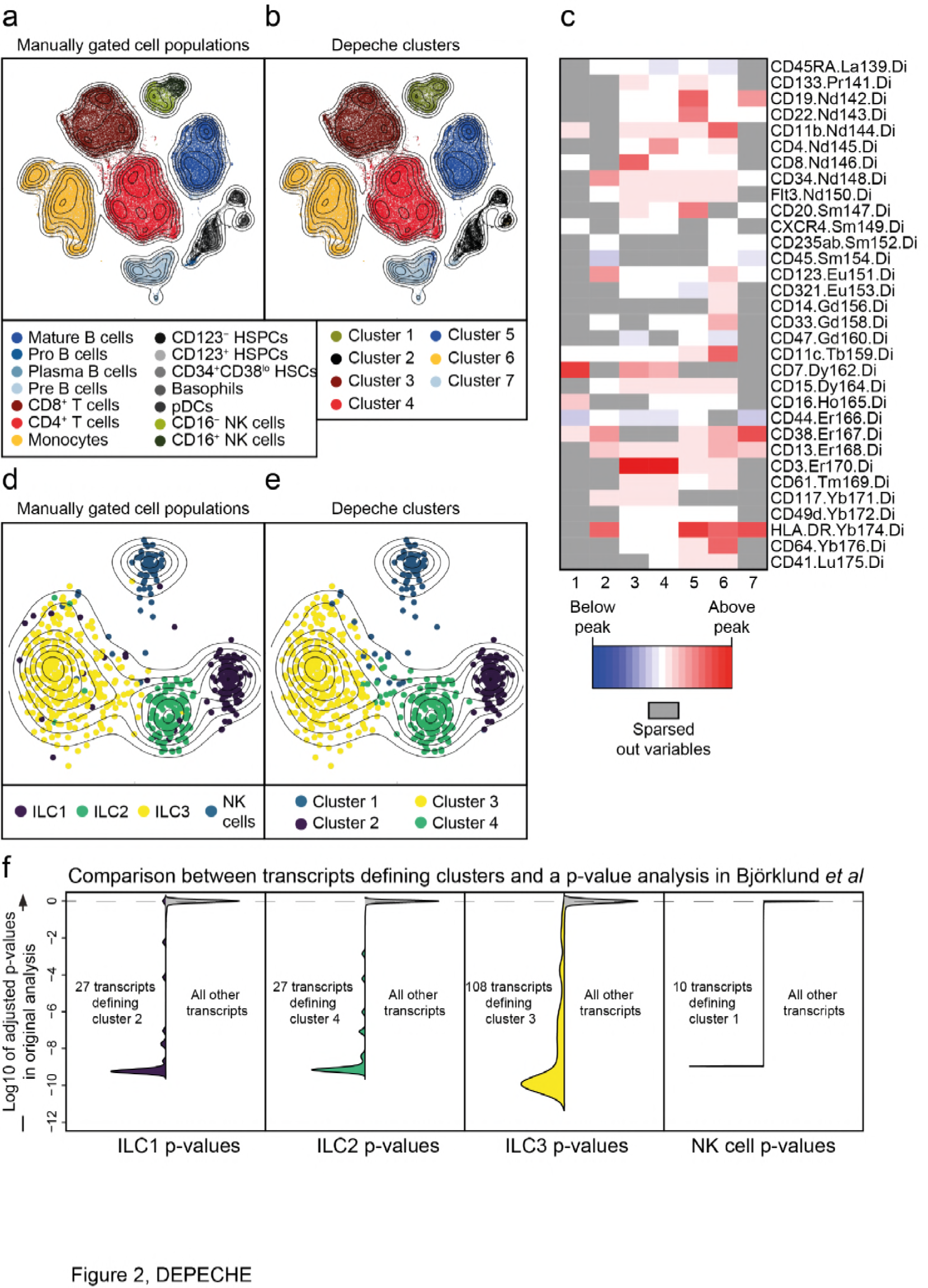
DEPECHE performance with real datasets with 32 or 35177 variables. a-b) bi-variate t-distributed stochastic neighbor embedding (tSNE) representation of the 32-variate mass cytometry data. A: distribution of manually defined cell populations over the tSNE field. B: distribution of DEPECHE clusters over the tSNE field. c) Heatmap showing which variables that define each cluster. Red color indicates a higher expression in the cluster than the most common expression for all observations. Blue color conversely indicates lower expression than the geometric mean for all observations. Grey color indicates that the variable in question does not contribute to defining the cluster. For Fig a-c all, 104184 cells have been clustered. d-tSNE representation of the 137-variate data subset that could efficiently distinguish the clusters in the 35177-variate single-cell transcriptome dataset. d) distribution of the cell types defined by index-sorting and manual gating on protein expression profiles shown over the tSNE field. e) distribution of DEPECHE clusters over the tSNE field. f) Violin plots illustrating the overlap between the original analysis by Björklund *et al* and the DEPECHE analysis. For each subplot, the left and right side illustrate the distribution of the transcripts defining the clusters, and all other transcripts, respectively. The y-axis shows the log10 of the p-values in the original analysis adjusted for multiple comparisons. For Fig d-f, all 648 cells have been clustered.

Single-cell transcriptomic datasets feature tens of thousands of variables. Thus, the need to exclude irrelevant variables is even more pressing, as compared to cytometry datasets. We therefore evaluated DEPECHE’s ability to cluster and extract the key transcripts defining clusters of a previously published single-cell transcriptomic dataset (Fig 2d-f) [19]. In this dataset, a total of 648 ILC1, ILC2, ILC3 and NK cells from three donors’ tonsils were index-sorted prior to RNA sequencing. Hence, these cell types, manually defined by protein expression, can be compared to clusters unbiasedly determined by RNA expression profiles [19]. In the DEPECHE analysis, no pre-selection of transcripts was performed, and hence, 35177 unique transcripts were included for each of the 648 cells. With the optimal penalty *λ*, four clusters were identified (Fig 2e). These corresponded well to the cell types as defined by protein expression; 84, 97, 91 and 97 percent of ILC1, ILC2, ILC3 and NK cells sorted into separate clusters, respectively (Fig 2d, e, S Fig 1b), leading to an average ARI of 0.78. Notably, cluster 1-4, corresponding to ILC1, ILC3, NK cells and ILC2, were defined by 27, 27, 108 and 10 transcripts, respectively (Fig 2f and Table 2), leading to a 99.9% average decrease in the number of variables. The transcripts identified to define the clusters in our analysis were among those most differentially expressed according to the original study [19] (Fig 2f). Thus, by identifying a finite number of variables, DEPECHE analysis increases interpretability and aides down-stream analyses. When DEPECHE clustering was compared to that of state-of-the-art algorithms [5–7] on the aforementioned dataset and six others (see Table 1), it performed consistently well as indicated by ARI (S Fig 2b). Thus, when applied to megavariate data, DEPECHE produces biologically relevant clusters and reduces the complexity of the result thousand-fold.

**Table 2:**
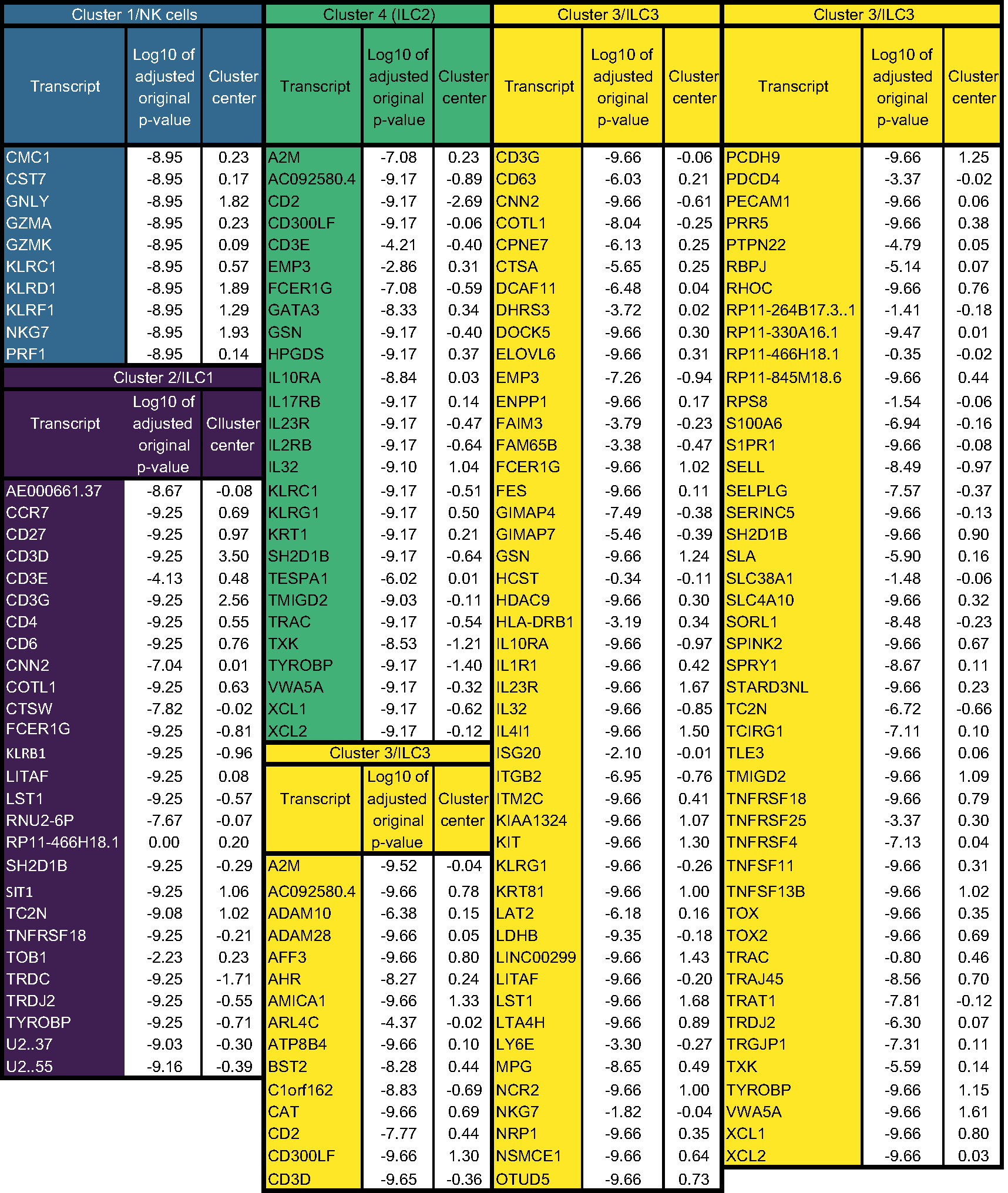
transcripts defining clusters in Björklund *et al* dataset

In conclusion, DEPECHE turns the penalized k-means methodology into a parameter free analysis technique guided by efficient calculation of the optimal clustering resolution. By doing so, it addresses the simultaneous problem of clustering and identification of biologically important variables that separate clusters. This is crucial in order to comprehend the noisy and often over-complicated data generated with current single cell technologies.

## Methods

### Clustering with DEPECHE

Clustering in DEPECHE is performed using a penalized version of the k-means algorithm, which is related to the k-means algorithm[8]. In this section, the k-means algorithm is outlined first, followed by an explanation of how it is extended to penalized k-means.

The k-means algorithm clusters data by fitting a mixture of normal distributions to the data with k equal mixture components and unit variance. Formally, *k d* -dimensional cluster centers, denoted *μ*_*i*,*j*_ where *i =* 1 … *k* and *j =* 1 … *d*, are fitted to the *n d* -dimensional datapoints *x*_*l*,*j*_, where *l =* 1 … *n*, by maximizing the score function

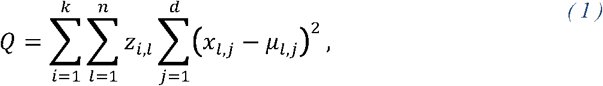

where *z*_*i*,*l*_ is 1 if the *l*th data point belongs to the *i*th cluster and zero otherwise. The score *Q* is optimized using an Expectation Maximization (EM) algorithm [20], *i.e.* so called E-and M-steps are iterated alternatingly until the score *Q* stops improving. In the E-step, the allocation variables *z*_*i*,*l*_ are updated so that each data point is allocated to its closest cluster. In the M-step, each cluster center *μ*_*i*,*j*_ is moved to the center of the data points allocated to it. When no more reallocation occurs in the E-step, the algorithm has converged.

In order to reduce the influence of uninformative dimensions that only contribute with noise, penalized k-means introduces an L1-penalty for each element of each cluster center *μ*_*i*,*j*_ to the optimization objective. Formally, the score function *Q* in Eq. (1), is updated:

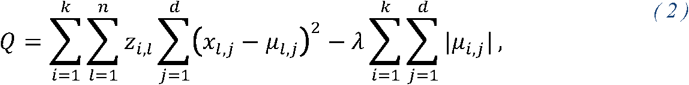

where *λ* is a positive penalty term. The additional term in the score function, introduced in Eq. (2) results in a change in the M-step of the original EM-algorithm of the k-means algorithm. Keeping *z*_*i*,*l*_ for all *l* fixed and optimizing *Q* with respect to *μ*_*i*,*j*_, the M-step is:

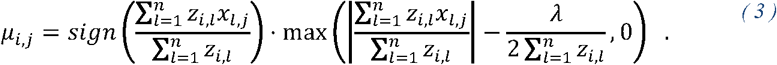

Bepending on the choice of the penalty parameter *λ*, some components of some clusters centers will be set to 0 in the M-step. Note that penalized k-means with penalty *λ =* 0 reduces to the original k-means algorithm.

In DEPECHE, cluster centers that are moved to the origin in the M-step are eliminated and not assigned any data points in the E-step. Due to the elimination of clusters, the number of produced clusters is independent of *k* and dependent on the penalty *λ* as long as at least one cluster is eliminated. In DEPECHE, *k* is always chosen to be so large that at least one cluster is eliminated.

Eq. (2) is a special case of the penalized model based clustering algorithm by Pan and Shen with unit variance and equal mixture components [9]. By imposing the penalty for each dimension and each cluster, penalized k-means identifies the dimensions that do not distinguish a particular cluster from the rest of the data, thus leaving these dimensions out of the definition of that cluster. This differs from the sparse k-means algorithm by Witten and Tibshirani [21] and the regularized k-means algorithm by Sun et al [10], that only identify dimensions that do not contribute to distinguish any cluster from the rest of the data.

Penalized k-means, as well as k-means, relies on a procedure for initializing the positions of the cluster centers. Cluster initialization is particularly delicate in DEPECHE, due to the elimination of clusters at the origin in the E-step. Poor initialization of the clusters might lead to elimination of too many clusters in the early E-steps, yielding fewer clusters in the end result than necessary to optimize *Q*. To avoid early elimination of clusters, DEPECHE initializes the cluster positions using the seed generation algorithm of k-means++ by Arthur and Vassilvitskii[22] and always starts clustering with penalty *λ =* 0. The penalty is then increased linearly over a number of E-steps until it reaches the predetermined value.

The EM-algorithm guarantees convergence to an optimum of the score *Q*, but not necessarily to the global optimum. In order to diminish the influence of the starting state, the EM-algorithm is run several times with random initialization, and the solution with optimal *Q* is chosen. In addition, *k* is set considerably higher than the expected number of final clusters, which also diminishes the sensitivity to the starting state. In the extreme case where *k* is set equal to the number of data points *n*, the outcome of penalized k-means is deterministic.

### Tuning the penalty

In this section, we describe the optimization scheme which is used for tuning the linear penalty *λ*. The outline of the algorithm:

1. Choose a range of penalty terms *λ*_*i*_, *i =* 1.. *N*_*λ*_ that are considered for clustering the dataset *D*
2. Create 2 datasets per penalty term *λ*_*i*_, called *D*_1,*i*_ and *D*_2,*i*_, by sampling *N*_*r*_ data points from *D* with replacement.
3. Run the penalized k-means algorithm on the datasets *D*_1,*i*_ and *D*_2,*i*_, yielding sets of cluster centers, denoted ?_1,*i*_ and ?_2,*i*_.
4. Create the partitions *P*_1,*i*_ and *P*_2,*i*_, by allocating all data points of the dataset *D* to their nearest cluster center of the sets ?_1,*i*_ and ?_2,*i*_.
5. Determine the Adjusted Rand Index (ARI), denoted *r* (*λ*_*i*_) from *P*_1,*i*_ to *P*_2,*i*_ [11].
6. Repeat step 2-5 times and average the obtained ARIs *r*(*λ*_*i*_) penalty wise until a stopping criteria regarding the statistical certainty of the obtained ARIs *r*(*λ*_*i*_) is met.
7. Choose the optimal penalty *λ*_*i*_, which is the penalty with the largest ARI *r*(*λ*_*i*_).

Some remarks to the parameter tuning procedure: The repetition Step 6 is necessary, since the obtained ARI *r*(*λ*_*i*_) is a random variable, due to the random procedure for creating the datasets *D*_1,*i*_ and *D*_2,*i*_ and the random procedure for initializing the penalized k-means algorithm. DEPECHE uses two stopping criteria: The first criterion creates an interval of width 2 standard errors around the obtained mean of *r*(*λ*_*i*_) and checks if the interval around the optimal ARI *r*(*λ*_*i*_) has a zero overlap with the other intervals. The second criterion checks whether the standard error of the mean of *r*(*λ*_*i*_) for the optimal penalty *λ*_*i*_ is below a threshold.

Step 2 requires a samples size *N*_*r*_. A natural choice is to set *N*_*r*_ equal to the number of data points, *n.* However, in cases where *n* is very large, so that the computational load of the optimization scheme becomes limiting, it is preferable to choose a smaller *N*_*r*_. In DepecheR, *N*_*r*_ *=* 10^4^ by default, in case *n* ≥ 10^4^. Notice that when an optimal penalty *λ*_*i*_ is discovered using sample size *N*_*r*_ ≠ *n*, the corresponding optimal penalty when sampling the full dataset *D* with magnitude *n* is (approximately) 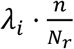, since the attraction force of a cluster is proportional to the number of data points in it. Exact calculation of the ARI in step 5 is computationally intractable for large datasets. Therefore, DEPECHE relies on an approximate ARI computation, based on 10^4^ random pairs of data points.

### Simultaneous Clustering and Parameter Tuning

For very large datasets (*n* > 10^8^), not only the penalty optimization, but also the final clustering once the optimal penalty has been found may be computationally intractable. However, increasing the size of the dataset, does not necessarily lead to an increase in number of clusters at the optimal resolution. In this case, it is feasible to cluster a subset of the full dataset *D* to obtain cluster centers *M* and then allocate the remaining data points of *D* to their closest clusters in *M*. This boosts computational efficiency since allocation imposes a much smaller computational load than clustering. Since several subsets of *D* are produced and clustered during the tuning of the penalty parameter *λ*, it seems natural to retrieve cluster centers *M* that were produced during the parameter tuning and use them to cluster *D*.

When picking a set of cluster centers *M* from the penalty tuning, the question arises which set of centers *M* to take, since several sets of centers, denoted *M*_*i*,*l*_ (*l =* 1,,, *p*), are produced for the optimal penalty *λ*_*i*_. In DEPECHE, the centers *M*_*i*,*j*_ that have the strongest similarity (on average) to the remaining *p* - 1 centers is chosen and is referred to as the most generalizable cluster set. The level of similarity between the centers *M*_*i*,*j*_ and *M*_*i*,*l*_ is quantified using the ARI between the partitions *P*_*i*,*j*_ and *P*_*i*,*l*_, induced by allocating each data point of *D* to its closest cluster center in *M*_*i*,*j*_ and *M*_*i*,*l*_ respectively.

#### Empiric Performance of the Penalty Tuning Scheme

Roughly speaking, DEPECHE combines a flavored penalized k-means algorithm with a parameter tuning scheme, which identifies an optimal resolution. A naturally arising question is then whether the parameter tuning scheme is able to determine a biologically relevant resolution or if other penalized k-means clustering resolutions outperform the resolution chosen by DEPECHE. Using a range of datasets (Table 3), the biological relevance (measured in ARI to the manually curated solution) of the optimized DEPECHE partitions were compared to the biologically optimal partition among all partitions generated with 20 repetitions on each of a range of 11 penalties per dataset. Overall, the DEPECHE resolution-selection showed close to optimal performance, as the selected solutions only had a median of 0.02 lower ARI to the gold standard (range 0-0.065) than the best possible solution with all penalties (Table 3).

**Table 3:** ARI between DEPECHE partitions and golden standard partitions

| Dataset | Median ARI in supplementary figure 2 | Maximal ARI with any penalty | Difference |
| --- | --- | --- | --- |
| Levine | 0.961 | 0.975 | 0.015 |
| Bendall | 0.841 | 0.873 | 0.032 |
| Biase | 1 | 1 | 0 |
| Björklund | 0.782 | 0.842 | 0.06 |
| Deng | 0.827 | 0.848 | 0.021 |
| Goolam | 0.629 | 0.639 | 0.009 |
| Kolodziejczyk | 0.992 | 1 | 0.008 |
| Pollen | 0.863 | 0.928 | 0.065 |
| Yan | 0.626 | 0.691 | 0.064 |

### Scaling and Centering the Data

The clusters produced by DEPECHE, as well as their interpretation, depends on the scaling and centering of the data. The scaling determines the relative importance of the measured variables, where variables with a larger spread have stronger influence on the clustering. The centering defines where zero occurs in each variable, thereby influencing the clustering results due to the linear penalty.

DEPECHE is applicable to a large range of datasets where the numbers of dimensions, *d*, and the number of data points, *n*, can vary with many orders of magnitude. The differing characteristics of these datasets require different treatments with respect to scaling and centering.

#### Scaling

Empirically, a majority of single-cell transcriptome datasets tend to have a few variables where the variance is many orders of magnitude greater than in the other variables. In this case, the high-variance variables will *de-facto* determine the clustering, implying that the clustering will fail to take the majority of the measured information into account. To even out the influence of these high-variance variables on the clustering outcome, the data is log transformed when such variables are present. In DepecheR, this data behavior is detected automatically by concatenating all variables into a one dimensional vector, for which the kurtosis is calculated. A high kurtosis, indicates that the variables differ greatly in their internal variance. For datasets with low kurtosis, refraining from the log transform is preferable, since transformation distorts the information.

#### Centering

Centering the origin to be close to the bulk of the data is preferable, in order to have all biological clusters at approximately the same distance from the origin. Having some biological clusters close to the origin and some far off is often unwanted, since the linear penalty then imposes a preference for creating clusters close to the origin. Apart from influencing the clustering, the centering also determines the interpretation of the obtained sparsity. Just as for scaling, which centering scheme to apply depends on the dataset.

For low dimensional datasets (*n*>100), DEPECHE applies maximal density centering, which sets the zero in each dimension to coincide with the highest data density. The density it computed by collecting the data in equally spaced bins (default number of bins in DepecheR is the number of data points *n* divided by 50), where the bin with the highest number of data points has the highest density. Using this scheme, sparsity (*i.e.* that a variable is non-contributing to the definition of a cluster) is interpreted as that the data points in the cluster do not deviate from the most common outcome. It also ensures that the origin is relatively close to the bulk of the data, since it is located at the most common outcome for each variable respectively. The benefit of this scheme is that it boosts sparsity, by declaring the most common outcome non-defining. However, for high dimensional datasets (*n* ≥ 100), maximal density centering can push the origin so far away from the center of mass of the dataset, that the penalty starts to impose an unwanted, artificial influence on the clustering, hampering the biological relevance of the clusters. To avoid this, DEPECHE imposes a mean centering scheme for such datasets, which locates the origin at the center of mass of the dataset.

A potential complication, related to centering, occurs when a biologically relevant cluster is located very close to the origin, since DEPECHE creates no clusters in the origin and will then force the cluster to merge with other clusters. However, this scenario was never detected in real data.

### Experimental procedures

#### Preprocessing of mass cytometry data

The benchmark datasets from Levine *et al* [12] and Bendall *et al*[18] were transformed using the flowTrans package [23] before used in any clustering algorithm.

#### Preprocessing of single-cell transcriptomic data

The dataset from Björklund *et al* [19] was normalized using the sva package [24] as in the original manuscript. For this dataset, doublet variables were removed, lowering the number of variables from 64443 to 35177.

The gold-standard datasets used for benchmarking in the publication by Kiselev *et al* [5] were obtained in a pre-processed state. Before clustering with any algorithm, the gene filter function used in the sc3 package was used [5], with settings removing the genes that were expressed in more than 90% of the cells. This resulted in the number of transcripts presented in Table 1 (range 13608-19571 transcripts).

### Code availability

All code necessary to generate the figures and tables in the manuscript are included in supporting code 1. The software package DepecheR is available for download at (https://github.com/theorell/depecher).

## Acknowledgments

The authors are grateful for all important input that has come from the initial users of the DepecheR software, especially Dr Niklas Björkström, Dr Jakob Michaëlsson and Sigrun Stultz. Other colleagues that have contributed seminally to the framework of ideas that have led to the creation of DEPECHE are Dr Bruce Bagwell and Dr Geoffrey Hart.

## Author contributions

A.T. drafted mathematical models, co-wrote the software implementation and the manuscript, Y.T.B. co-wrote the manuscript, J.T. co-wrote mathematical models and drafted software implementation and the manuscript.

## Supporting information

**Fig S1:**
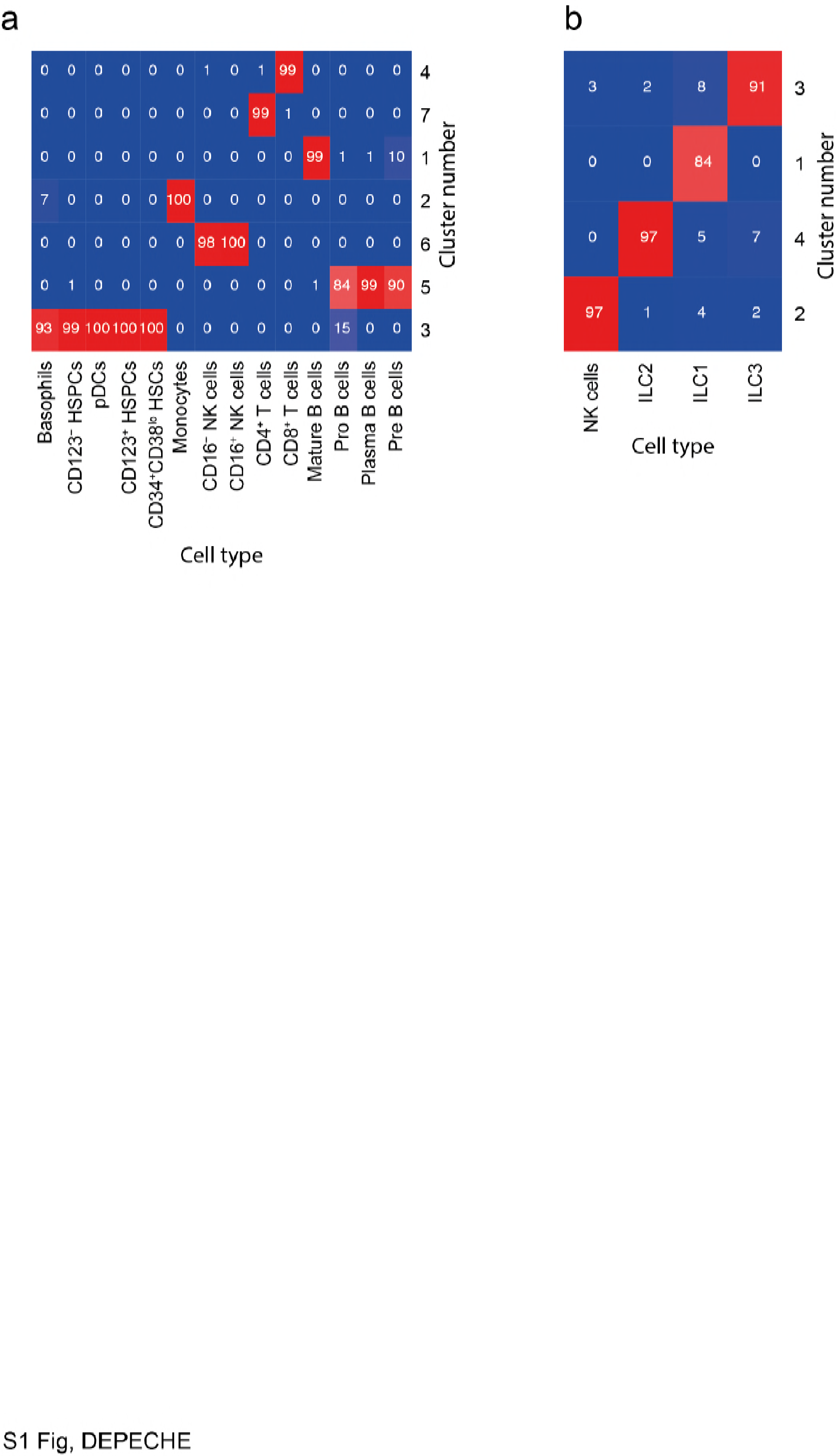
Heatmaps comparing the golden standard partitions to the DEPECHE partitions for a) the 32-variate Levine dataset and b) the 35166-variate Björklund dataset. Red color indicates large overlap, blue color indicates low overlap between a gold standard-vs-depeche cluster pair.

**Fig S1:**
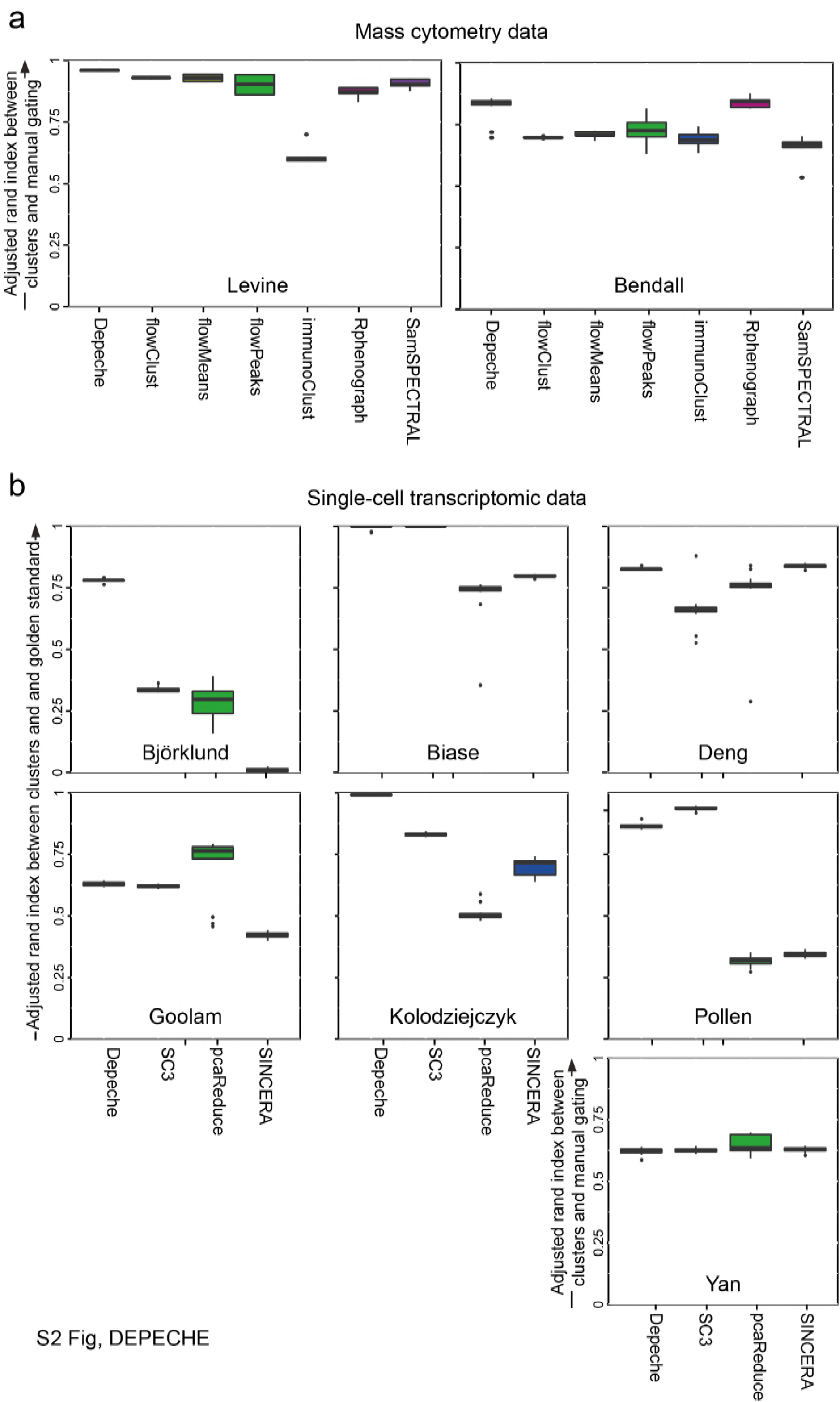
Algorithm comparisons. For all graphs, the x-axis shows the algorithms and the y-axis shows the Adjusted Rand Index comparing the clustering result with the golden standard clustering. a) Subsamples with 20000 unique cells from two mass cytometry datasets published by Levine *et al* and Bendall *et al* were clustered with DEPECHE and six previously published algorithms. For each dataset and algorithm, clustering was performed on 20 unique subsamples. For flowClust, flowPeaks and SamSPECTRAL, that do not perform internal parameter tuning, a range of parameter values were evaluated and the parameter value sets generating the highest ARI values were selected for display. b) The full Björklund dataset, as well as six other datasets previously used for benchmarking by Kiselev *et al* were clustered 20 times with DEPECHE and three other algorithms. The Björklund dataset was normalized to reduce batch effects, with the procedure described in the original publication. These six datasets were also automatically log2-transformed within DEPECHE, and thus, log2-transformation was applied also for Sincera and pcaReduce, whereas sc3 was fed both log2-and untransformed data. The lower and upper hinges of all boxplots extend to the 25:th and 75:th percentile, whereas the line in the middle describes the median. The whiskers extend to the lowest and highest value no further than 1.5 times the distance between the 25:th and 75:th percentile. Outside of this range, the observations are considered outliers and are shown as dots.

**S1 File. The DepecheR software, for the review phase.**

**S2 File. The code needed to generate all figures, for the review phase.**

Author contributions
A.T. drafted mathematical models, <u>coded</u> the software implementation and co-wrote the manuscript, Y.T.B. co-wrote the manuscript, J.T. drafted mathematical models, coded software implementation and wrote the manuscript.

## References

1. Spitzer MH, Nolan GP. Mass Cytometry: Single Cells, Many Features. Cell. 2016;165: 780–791. doi:10.1016/j.cell.2016.04.019

2. Tanay A, Regev A. Scaling single-cell genomics from phenomenology to mechanism. Nature. 2017;541: 331–338. doi:10.1038/nature21350

3. Budnik B, Levy E, Slavov N. Mass-spectrometry of single mammalian cells quantifies proteome heterogeneity during cell differentiation. bioRxiv. 2017; 102681. doi:10.1101/102681

4. Saeys Y, Gassen SV, Lambrecht BN. Computational flow cytometry: helping to make sense of high-dimensional immunology data. Nat Rev Immunol. 2016;16: 449–462. doi:10.1038/nri.2016.56

5. Kiselev VY, Kirschner K, Schaub MT, Andrews T, Yiu A, Chandra T, et al. SC3: consensus clustering of single-cell RNA-seq data. Nat Methods. 2017;14: 483–486. doi:10.1038/nmeth.4236

6. Guo M, Wang H, Potter SS, Whitsett JA, Xu Y. SINCERA: A Pipeline for Single-Cell RNA-Seq Profiling Analysis. PLOS Comput Biol. 2015;11: e1004575. doi:10.1371/journal.pcbi.1004575

7. Žurauskienė J, Yau C. pcaReduce: hierarchical clustering of single cell transcriptional profiles. BMC Bioinformatics. 2016;17: 140. doi:10.1186/s12859-016-0984-y

8. MacQueen J. Some methods for classification and analysis of multivariate observations. Proceedings of the Fifth Berkeley Symposium on Mathematical Statistics and Probability. Berkeley, California: University of California Press; 1967. pp. 281–297. Available: https://projecteuclid.org/euclid.bsmsp/1200512992

9. Pan W. Penalized model-based clustering with application to variable selection. J Mach Learn Res. 2007;8: 1145–1164.

10. Sun W, Wang J, Fang Y. Regularized k-means clustering of high-dimensional data and its asymptotic consistency. Electron J Stat. 2012;6: 148–167. doi:10.1214/12-EJS668

11. Hubert L, Arabie P. Comparing partitions. J Classif. 1985;2: 193–218. doi:10.1007/BF01908075

12. Levine JH, Simonds EF, Bendall SC, Davis KL, Amir ED, Tadmor MD, et al. Data-Driven Phenotypic Dissection of AML Reveals Progenitor-like Cells that Correlate with Prognosis. Cell. 2015;162: 184–197. doi:10.1016/j.cell.2015.05.047

13. Lo K, Hahne F, Brinkman RR, Gottardo R. flowClust: a Bioconductor package for automated gating of flow cytometry data. BMC Bioinformatics. 2009;10: 145. doi:10.1186/1471-2105-10-145

14. Aghaeepour N. flowMeans: Non-parametric Flow Cytometry Data Gating. 2010.

15. Sörensen T, Baumgart S, Durek P, Grützkau A, Häupl T. immunoClust-An automated analysis pipeline for the identification of immunophenotypic signatures in high-dimensional cytometric datasets. Cytom Part J Int Soc Anal Cytol. 2015;87: 603–615. doi:10.1002/cyto.a.22626

16. Ge Y, Sealfon SC. flowPeaks: a fast unsupervised clustering for flow cytometry data via K-means and density peak finding. Bioinforma Oxf Engl. 2012;28: 2052–2058. doi:10.1093/bioinformatics/bts300

17. Zare H, Shooshtari P, Gupta A, Brinkman RR. Data reduction for spectral clustering to analyze high throughput flow cytometry data. BMC Bioinformatics. 2010;11: 403. doi:10.1186/1471-2105-11-403

18. Bendall SC, Simonds EF, Qiu P, Amir ED, Krutzik PO, Finck R, et al. Single-Cell Mass Cytometry of Differential Immune and Drug Responses Across a Human Hematopoietic Continuum. Science. 2011;332: 687–696. doi:10.1126/science.1198704

19. Björklund ÅK, Forkel M, Picelli S, Konya V, Theorell J, Friberg D, et al. The heterogeneity of human CD127(+) innate lymphoid cells revealed by single-cell RNA sequencing. Nat Immunol. 2016;17: 451–460. doi:10.1038/ni.3368

20. Dempster AP, Laird NM, Rubin DB. Maximum Likelihood from Incomplete Data via the EM Algorithm. J R Stat Soc Ser B Methodol. 1977;39: 1–38.

21. Witten DM, Tibshirani R. A framework for feature selection in clustering. J Am Stat Assoc. 2010;105: 713–726. doi:10.1198/jasa.2010.tm09415

22. Arthur D, Vassilvitskii S. K-means++: The Advantages of Careful Seeding. Proceedings of the Eighteenth Annual ACM-SIAM Symposium on Discrete Algorithms. Philadelphia, PA, USA: Society for Industrial and Applied Mathematics; 2007. pp. 1027–1035. Available: http://dl.acm.org/citation.cfm?id=1283383.1283494

23. Finak G, Manuel-Perez J, Gottardo R. flowTrans: Parameter Optimization for Flow Cytometry Data Transformation. 2010.

24. Leek JT, Johnson WE, Parker HS, Jaffe AE, Storey JD. The sva package for removing batch effects and other unwanted variation in high-throughput experiments. Bioinforma Oxf Engl. 2012;28: 882–883. doi:10.1093/bioinformatics/bts034

